# The Endothelial Protein C Receptor plays an essential role in the maintenance of Pregnancy

**DOI:** 10.1101/2020.02.05.935940

**Authors:** Michelle M Castillo, Qiuhui Yang, Abril Solis Sigala, Dosia T McKinney, Min Zhan, Kristen L Chen, Jason A Jarzembowski, Rashmi Sood

## Abstract

Placenta-mediated pregnancy complications are a major challenge in the management of maternal-fetal health. Maternal thrombophilia is a suspected risk factor but the role of thrombotic processes in these complications and the potential for antithrombotic treatment have remained unclear. Endothelial Protein C Receptor (EPCR) is an anticoagulant protein highly expressed in the placenta. EPCR autoantibodies and specific gene variants of EPCR are associated with poor pregnancy outcomes. In mice, fetal EPCR deficiency results in placental failure and *in utero* death. Adult EPCR-deficient mice generated by maintaining placental expression exhibit plasma markers of thrombophilia without overt thrombosis. We demonstrate that inactivation of clotting factor VIII or Protease Activated Receptor 4 (Par4), Par3 or integrin αIIb in the mother allows placental development and intrauterine survival of murine embryos lacking EPCR. Rescued EPCR-deficient embryos exhibit thrombosis in placental venous sinuses at late gestation and a high rate of neonatal lethality. In contrast to fetal EPCR deficiency, maternal deficiency of EPCR results in frequent stillbirths and maternal death accompanied by pathological findings that resemble placental abruption and consumptive coagulopathy. Inactivation of Par4, but not clotting factor VIII, prevents maternal death and restores normal pregnancy outcomes. These observations establish a cause-effect relationship between maternal thrombophilia and placental abruption. They demonstrate that sites of uteroplacental thrombosis and the potential response to antithrombotic intervention may differ with gestational age and maternal versus fetal origin of thrombophilia. Our findings highlight the potential for therapeutic inhibition of thrombin-mediated platelet activation in a subset of pregnancy complications.

**KEY POINTS:**

1. Murine model establishes a cause-effect relationship between maternal thrombophilia, retroplacental hemorrhage and severe pregnancy complications.
2. Thrombin-mediated activation of maternal platelets is a key event in thrombophilia-associated pregnancy complications and a potential target of therapeutic intervention.
3. Maternal venous channels in uteroplacental circulation are additional sites of thrombotic pathology associated with adverse neonatal outcomes.

## INTRODUCTION

Thrombophilia is a suspected risk factor for placenta-mediated pregnancy complications that include recurrent pregnancy loss, intrauterine growth restriction, stillbirth, placental abruption and preeclampsia ^1,2^. These complications affect a significant proportion of all pregnancies and are a major cause of maternal and fetal morbidity and mortality ^3-5^. Treatment of these disorders with low molecular weight heparin (LMWH) has shown mixed outcomes, with recent meta-analyses suggesting absence of significant benefit ^6-8^. Humans and rodents form a hemochorial placenta where extraembryonic trophoblast cells replace maternal vascular endothelium. The interfaces of maternal blood and trophoblast cells are potential sites of pathologic coagulation ^9^. At these sites, maternal platelets could participate in thrombosis and/or engage inflammatory mechanisms causing pregnancy complications. Given the low 50% effective concentration (EC_50_) of thrombin-mediated platelet activation, it is unclear if treatment with LMWH reduces local thrombin generation to levels below those needed for platelet activation. In this work, we first used murine models to rigorously examine whether maternal platelet activation plays a key role in placental failure due to fetal thrombophilia. We next investigated if maternal thrombophilia is associated with pregnancy complications and the potential for anti-platelet and anti-thrombotic therapy.

The endothelial protein C receptor (EPCR) is an anticoagulant protein highly expressed on trophoblast cells ^9,10^. EPCR binds to protein C (PC) and facilitates its activation by the thrombin-thrombomodulin complex. Activated PC (aPC) inhibits excessive thrombin generation by limited proteolytic cleavage and inactivation of coagulation factors Va and VIIIa. EPCR gene polymorphisms and presence of EPCR autoantibodies in the mother are associated with recurrent miscarriages and unexplained pregnancy loss ^11-14^. In mice, absence of EPCR in the fetus results in placental thrombosis and mid-gestational fetal death ^15^. Intrauterine death of EPCR-deficient mice is prevented by concomitant reduction in tissue factor (TF) or by maintaining EPCR expression on trophoblast cells. Yet, treatment of pregnant dams with LMWH prolongs survival of only a small fraction of EPCR-deficient embryos ^16^. Mice carrying a variant of EPCR with impaired ability to bind PC/aPC are viable without reported placental thrombosis and intrauterine death ^17^. Though most well studied for their role in anticoagulation, new roles of aPC and EPCR in stem cell function ^18-22^, inflammation ^23-25^ and immunity ^26,27^ have recently emerged. Thus, the mechanism of placental failure and whether it is primarily mediated by the loss of anticoagulation by EPCR remains unclear.

## RESULTS

### Thrombin activation of maternal platelets kills EPCR-deficient embryos

Platelet activation is an integral part of thrombosis. Maternal platelets surround the implantation site in close proximity to extraembryonic trophoblast cells in murine (Figure 1) and human placenta ^28^. Their activation could contribute to placental failure of EPCR-deficient embryos. Analogous to Protease Activated Receptor (PAR) 1 and PAR4 on human platelets, murine platelets express dual thrombin receptors Par3 and Par4. Par3 promotes cleavage and activation of Par4 at low thrombin concentration. In the absence of Par3, platelets continue to be activated via Par4 but the EC_50_ of platelet activation by thrombin increases from 0.10 ± 0.08 nM to 1.5 ± 0.7 nM ^29^. Murine platelets lacking Par4 are unresponsive to thrombin ^30^. We conducted genetic studies to determine if activation of maternal platelets via thrombin receptor Par3 and Par4 plays a critical role in placental failure of EPCR-deficient mice. Mice lacking EPCR die *in utero* before 11.5 days post coitum (dpc) in pregnancies from EPCR^+/-^ intercrosses (Table 1, rows 1 & 2; and reported by Gu *et. al.* ^15^). We analyzed pregnancies of Par4^-/-^ EPCR^+/-^ dams sired by EPCR^+/-^ males at 11.5 dpc (Table 1, row 3). Absence of Par4 in the mother allowed embryonic development of EPCR^-/-^ embryos (Table 1, compare rows 1 & 3, P<<0.001, χ^2^ test of independence). These were normal in appearance and showed placental development comparable to littermate EPCR^+/+^ controls (Figure 2 A-F). In this genetic experiment, the mother lacked Par4 and the fetuses (and their extraembryonic tissues) were heterozygous for Par4. Par4 is expressed on a variety of cell types that include trophoblast cells ^9^. To confirm that the observed effects reflect maternal Par4 deficiency rather than fetal haploinsufficiency, we performed reverse genetic crosses where Par4^-/-^ EPCR^+/-^ males sired pregnancies of EPCR^+/-^ dams. Absence of Par4 in the father alone did not protect EPCR^-/-^ embryos from mid-gestational death; no live Par4^+/-^ EPCR^-/-^ embryos were found at 11.5 dpc in these pregnancies (Table 1, row 4). Thus, maternal, but not paternal, deficiency of Par4 protects EPCR-deficient embryos from placental failure and mid-gestational demise. We next analyzed pregnancies of Par3^-/-^ EPCR^+/-^ dams sired by EPCR^+/-^ males. The absence of Par3 in the mother restored Mendelian frequency of EPCR-deficient embryos at 11.5 dpc (Table 1, row 5) and conferred significant protection (Table 1, compare rows 1 & 5, P<<0.001, χ^2^ test of independence). EPCR^-/-^ placentae appeared normal and unremarkable upon histological examination (Figure 2 G-L). Reverse genetic crosses (pregnancies of EPCR^+/-^ females sired by Par3^-/-^ EPCR^+/-^ males) did not produce any live EPCR^-/-^ embryos at 11.5 dpc and exhibited a high abortion rate (7 EPCR^+/+^, 7 EPCR^+/-^, 0 EPCR^-/-^, 10 aborted, 3 pregnancies analyzed, P=0.030, χ^2^ GOF). These results strongly suggested that thrombin-mediated activation of maternal platelets plays a key role in placental failure of EPCR-deficient embryos.

**Table 1.**
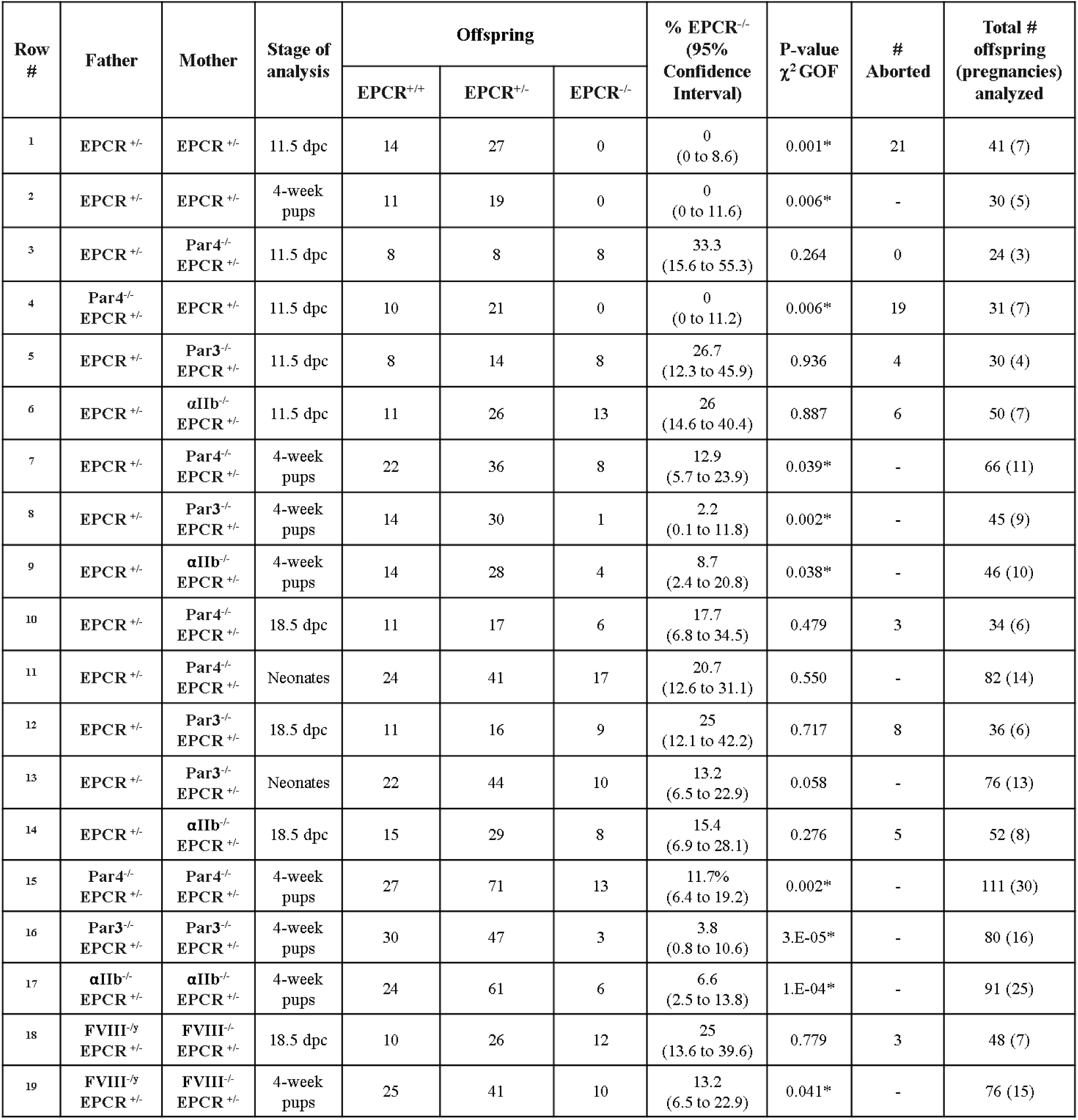
Pregnancy outcomes from EPCR^+/-^ intercrosses with and without inactivation of Par4, Par3, αIIb or FVIII in the mother, father or both parents.

**Figure 1:**
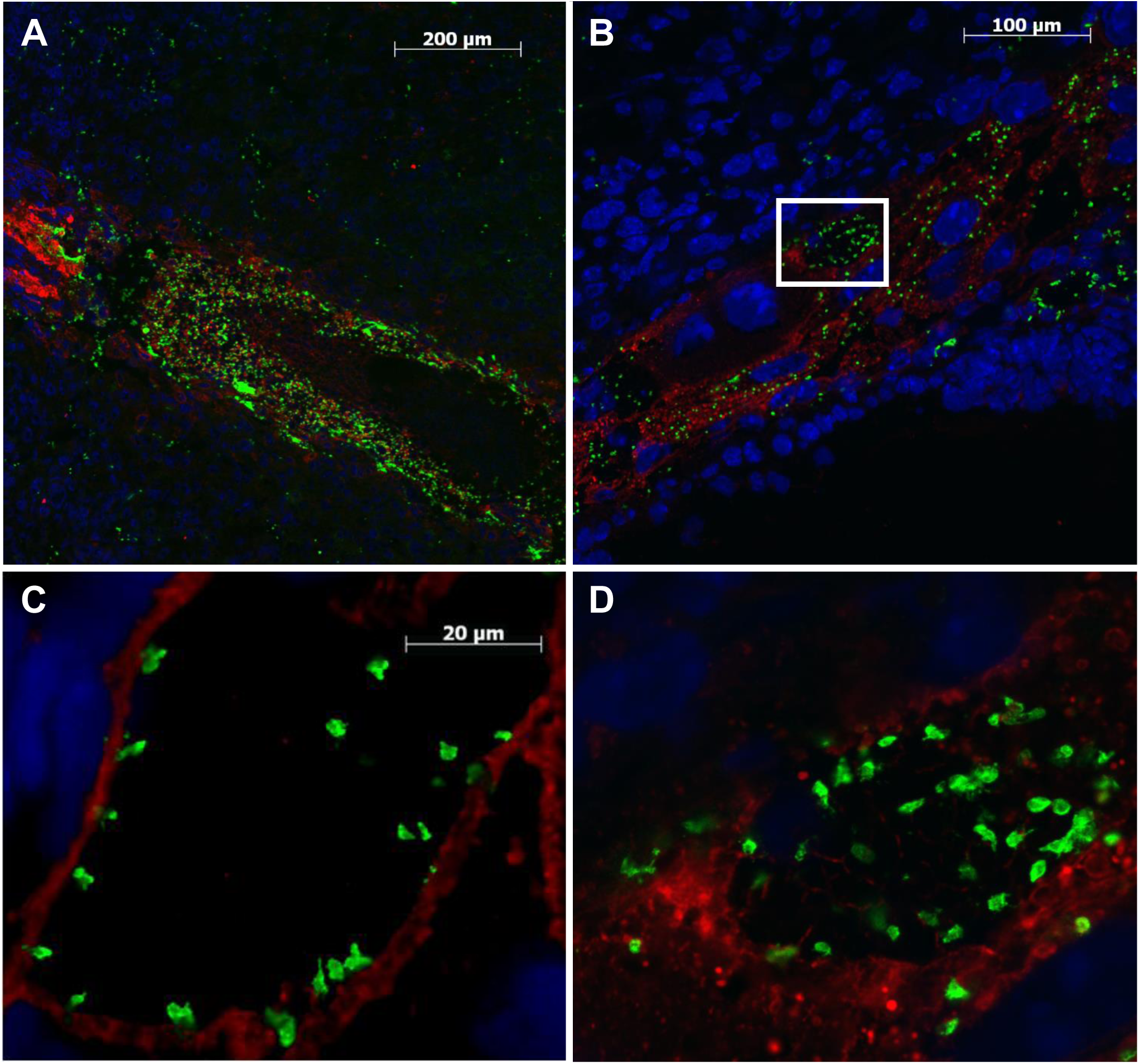
Platelets (stained green with GP1bβ antibodies) surround the implantation site of murine placenta at 7.5 dpc (**A**) and are seen in close proximity to trophoblast cells (stained red with cytokeratin antibodies) throughout early placental development (**B** & **D** 9.5 dpc, **C** 10.5 dpc). Nuclei are stained blue with DAPI. D is a magnified image of the boxed area in B. Scale bars represent 200µm (A), 100µm (B) and 20µm (C & D).

**Figure 2:**
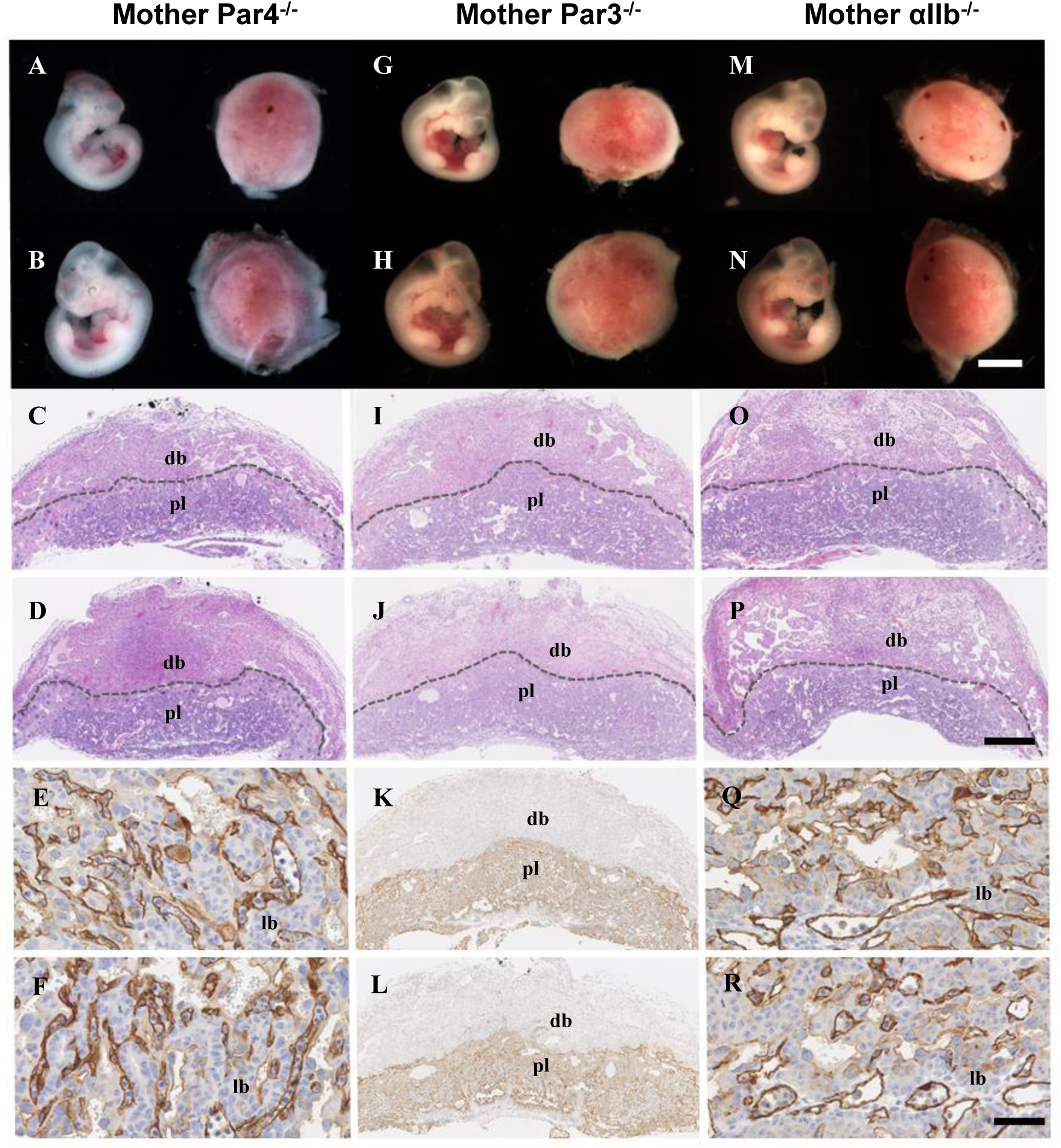
Genetic inactivation of Par4, Par3 or αIIb in the mother allows placental and embryonic development of EPCR-deficient mice past 10.5 dpc. Representative images of progeny analyzed at 11.5 dpc in pregnancies of Par4^-/-^ EPCR^+/-^ (**A-F**), Par3^-/-^ EPCR^+/-^ (**G-L**) or αIIb^-/-^ EPCR^+/-^ (**M-R**) dams sired by EPCR^+/-^ males are shown. Top two rows show whole mounts of EPCR^-/-^ (A, G, M) and littermate EPCR^+/+^ (B, H, N) embryos and placentae. Middle two rows show hematoxylin-eosin stained histological sections of placentae corresponding to EPCR^-/-^ (C, I, O) and EPCR^+/+^ (D, J, P) embryos. Dashed lines delineate placenta (pl) from decidua basalis (db). Bottom two rows show amplified images of labyrinth (lb) regions of EPCR^-/-^ (E and Q) and EPCR^+/+^ (F and R) placentae immunostained with CD31 antibodies or images of EPCR^-/-^ (K) or EPCR^+/+^ (L) placentae immunostained with cytokeratin antibodies. Scale bars represent 2mm (whole mounts) or 500μm (placental sections).

Aspirin reduces platelet aggregation by inhibiting Cox-1 and Cox-2 enzymes and has been tested for its ability to improve outcomes in human pregnancy. In EPCR^+/-^ intercrosses, aspirin treatment of pregnant dams prolonged the survival of EPCR-deficient embryos but did not restore normal placental development (Supplementary note 1 and Figure S1 A-E). In contrast, depletion of maternal platelets restored placental development and allowed survival of EPCR-deficient embryos (Supplementary note 2 and Figure S1 F-I). Upon activation by thrombin, platelets convert integrin αIIbβ3 to its high affinity state. Integrin αIIbβ3 mediates binding to fibrinogen and von Willebrand factor and plays a key role in platelet aggregation. Inhibitors of αIIbβ3 that block this common pathway of platelet adhesion and aggregation have been used in the management of acute coronary syndromes ^31^. We tested the role of integrin αIIbβ3 in placental failure of EPCR-deficient mice. Genetic inactivation of αIIb in the mother fully protected EPCR^-/-^ embryos from mid-gestational death (Table 1, row 6). On histological examination, αIIb^+/-^ EPCR^-/-^ placentae from these crosses were unremarkable at 11.5 dpc (Figure 2 M-R). In contrast, EPCR-deficient embryos were not found in pregnancies of EPCR^+/-^ dams sired by αIIb^-/-^ EPCR^+/-^ males. Rather, we observed a high rate of abortions (6 EPCR^+/+^, 14 EPCR^+/-^, 0 EPCR^-/-^, 7 aborted, 4 pregnancies analyzed at 18.5 dpc). Thus, inactivation of Par4, Par3 or the integrin αIIbβ3 receptor on maternal platelets prevented placental failure and mid-gestational death of EPCR-deficient embryos, but aspirin treatment provided marginal benefit. Taken together, these data demonstrate that thrombin-mediated activation of maternal platelets causes placental failure and mid-gestational death of EPCR-deficient embryos.

### Placental thrombosis leads to high neonatal lethality of EPCR-deficient pups

Although the above genetic approaches allowed placental and embryonic development of EPCR-deficient mice past mid-gestation, we observed that they were present at significantly reduced frequencies among 4-week old pups genotyped at wean (Table 1, rows 7, 8 & 9). This was a surprising finding as maintaining EPCR expression on trophoblast cells and specifically deleting it from epiblast derivatives results in the generation of mice with undetectable level of EPCR expression. EPCR-deficient mice generated by this approach are born at expected Mendelian frequency and exhibit normal life spans ^16^. We independently repeated these results in our laboratory. In pregnancies of EPCR^lox/lox^ females sired by Meox2Cre^tg/-^ EPCR^δ/δ^ males, we observed 27 Meox2Cre^-/-^ EPCR^lox/δ^ and 23 Meox2Cre^tg/-^ EPCR ^δ/δ^ live pups at 4 weeks of age (P= 0.85, χ2 GOF test; 50% EPCR^-/-^ expected, 46.0% observed, 95% CI: 31.8 to 60.7%). These results confirm that epiblast-specific deletion of EPCR (Meox2Cre^tg/-^ EPCR^δ/δ^; abbreviated henceforth as EPCR^δ/δ^) allows embryonic development and survival comparable to EPCR-expressing littermate controls. Thus, though the absence of Par4, Par3 or αIIb in the mother protected EPCR-deficient (EPCR^-/-^) embryos from mid-gestational death, longer-term survival of these mice that lack EPCR expression on extraembryonic trophoblast cells was markedly reduced, compared to the survival of EPCR-deficient mice generated by maintaining expression on trophoblast cells (EPCR^δ/δ^).

To investigate the time and cause of death of EPCR^-/-^ embryos/pups, we examined pregnancy outcomes at 18.5 dpc and within the first 24 hours of birth. Frequencies of late gestation EPCR^-/-^ embryos and neonates from pregnancies of Par4^-/-^ EPCR^+/-^ dams sired by EPCR^+/-^ males did not deviate significantly from expected (Table 1, rows 10 & 11). EPCR^-/-^ offspring were also present in expected Mendelian frequency at 18.5 dpc in pregnancies of Par3^-/-^ EPCR^+/-^ dams sired by EPCR^+/-^ males (Table 1, row 12) but were markedly reduced among neonates (Table 1, row 13, P=0.058, χ^2^ GOF). Fewer than expected EPCR-deficient mice were found at late gestation in pregnancies of αIIb ^-/-^ EPCR^+/-^ mothers sired by EPCR^+/-^ males, but the reduction in their frequency did not reach statistical significance (Table 1, row 14, 15.5% observed, 25% expected, P=0.276, χ^2^ GOF). In each of these crosses, 100% of EPCR^-/-^ mice that survived past the first week of birth exhibited normal life spans (> 1 year of age). Thus, EPCR-deficient pups are vulnerable in the perinatal period, and many died within the first few days of birth. We examined EPCR^-/-^ neonates from each of these genetic crosses but did not discover any obvious potential cause of death. However, we observed that EPCR-deficient embryos in pregnancies of Par3^-/-^ EPCR^+/-^ dams were significantly smaller than their littermates at 18.5 dpc. The reduction in size was also observed for their corresponding placentae. A similar reduction was noted in placental sizes for EPCR-deficient concepti in pregnancies of Par4^-/-^ EPCR^+/-^ and αIIb^-/-^ EPCR^+/-^ dams and in sizes of EPCR-deficient embryos in pregnancies of αIIb^-/-^ EPCR^+/-^ dams (Figure 3 A-J). Histological examination of late gestation placentae in each of these genetic crosses strikingly revealed thrombi in the junctional zone of EPCR^-/-^ placentae. EPCR is highly expressed in the spongiotrophoblast cells of the junctional zone in normal mouse placentae. The junctional zone is traversed by trophoblast lined venous sinuses that drain maternal blood from the labyrinth to decidua basalis ^32^. Frequent thrombi were observed in these channels in EPCR^-/-^ but not in littermate EPCR^+/+^ placentae (Figure 3 K-T). Calcified and infarcted regions were also frequently observed in the placental labyrinth, but these did not correlate with any specific genotype. EPCR^-/-^ mice continued to exhibit neonatal lethality even when both maternal and fetal genes for Par4, Par3 or αIIb were inactivated (Table 1, rows 15, 16 & 17). We examined 18.5 dpc EPCR^-/-^ placentae among progeny of αIIb^-/-^ EPCR^+/-^ intercrosses. Thrombi persisted in the junctional zone of these placentae (Figure 3 U & V). In contrast, thrombi were not present in the junctional zone of placentae corresponding to Meox2Cre^tg/-^ EPCR^-/-^ embryos where extraembryonic EPCR expression was maintained on spongiotrophoblast cells (Figure 3 W-Z). These data show a remarkable association between placental thrombosis (due to EPCR deficiency on trophoblast cells) and the high neonatal lethality of EPCR^-/-^ mice. Neonatal lethality is not observed when EPCR expression is maintained on trophoblast cells but removed from the embryo. Inactivation of maternal Par4, Par3 or αIIb each improved survival, but did not completely overcome placental thrombosis and associated neonatal death.

**Figure 3:**
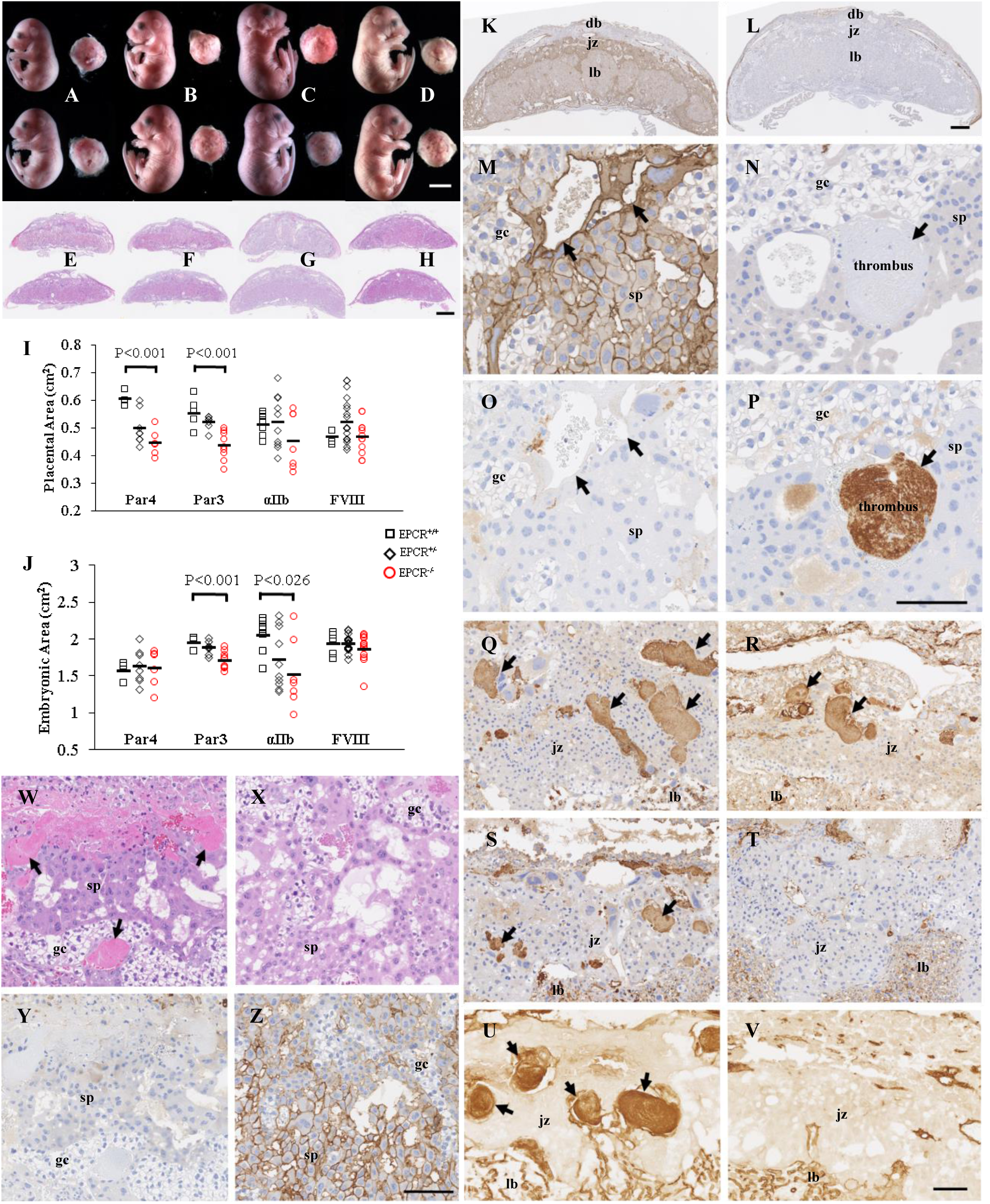
EPCR-deficient embryos survive to late gestation, but placentae show thrombi in venous sinuses of the junctional zone. **A-D**: Whole mounts of EPCR^-/-^ embryos and placentae (top row) and littermate EPCR^+/+^ controls (bottom row) from pregnancies of Par4^-/-^ EPCR^+/-^ (A, E), Par3^-/-^ EPCR^+/-^ (B, F) αIIb^-/-^ EPCR^+/-^ (C, G) or FVIII^-/-^ EPCR^+/-^ (D, H) dams. **E-H**: Hematoxylin-eosin stained histological sections of placentae corresponding to A-D (top row EPCR^-/-^, bottom row EPCR^+/+^). **I-J**: Cross-sectional area of the placental disc (I) and lengthwise cross-sectional area of the embryos (J) measured from images of whole mounts, grouped by maternal genes that were inactivated. **K-P**: Histological sections of EPCR^+/+^ (K, M and O) and EPCR^-/-^ (L, N, and P) placentae from pregnancies of αIIb^-/-^ EPCR^+/-^ dams sired by EPCR^+/-^ males at 18.5 dpc. M and N are enlarged images of the junctional zones from K and L respectively. K-N are immunostained with EPCR antibodies and O-P with fibrin(ogen) antibodies. Black arrows point to venous channels that traverse the junctional zone. Thrombi are seen in the venous channels in EPCR^-/-^ placentae (black arrows in N and P). **Q-V**: Thrombi immunostained with anti-fibrin(ogen) antibodies are also seen in the junctional zones of EPCR^-/-^ placentae from pregnancies of Par4^-/-^ EPCR^+/-^ (Q), or Par3^-/-^ EPCR^+/-^ (R) sired by EPCR^+/-^ males or FVIII^-/-^ EPCR^+/-^ intercrosses (S), but not in the placentae of EPCR^+/+^ controls (T). Venous thrombi persist in EPCR^-/-^ placentae (U) but are not seen in EPCR^+/+^ placentae (V) from pregnancies of αIIb^-/-^ EPCR^+/-^ intercrosses. U-V are immunostained with PECAM (CD31) antibodies that recognize platelets and endothelial cells. **W-Z**: Thrombi are seen in the junctional layer of 15.5 dpc placentae that lack EPCR expression (pregnancies of Par4^-/-^ EPCR^+/-^ dams; W, Y) but not in placentae of EPCR^δ/δ^ embryos that express EPCR on trophoblast cells (X). Y and Z are EPCR immunostained sections adjacent to hematoxylin-eosin stained sections shown in W and X, respectively. Scale bars represent 5mm (A-D), 1mm (E-H), 500µm (K, L) and 100µm (M-P, Q-V, W-Z). db: Decidua basalis; jz: junctional zone; lb: labyrinth; sp: spongiotrophoblasts; gc: glycogen cells.

### Inactivation of coagulation factor VIII protects EPCR-deficient embryos

Daily injections of 20μg/g of LMWH was reported to prolong survival of a small fraction of EPCR-deficient mice; 2 out of 48 pups born from heterozygous intercrosses were found to be EPCR^-/-^ (4.2% observed, 25% expected) ^16^. In our laboratory, continuous delivery of LMWH through subcutaneously implanted osmotic pumps also resulted in marked anticoagulation but did not restore normal placental development (Supplementary note 3 and Figure S2). To rigorously test the role of excess thrombin generation in fetal demise, we conducted breeding experiments using mice in which FVIII was genetically inactivated. FVIIIa supports the propagation of thrombin generation via intrinsic tenase (FVIIIa-FIXa) formation. It is one of the targets of inactivation by aPC. We analyzed pregnancies of FVIII^-/-^ EPCR^+/-^ dams sired by FVIII^-/y^ EPCR^+/-^ males. (Note that FVIII is an X-linked gene with a single copy in males.) EPCR^-/-^ embryos were present at expected Mendelian frequency at 18.5 dpc (Table 1, row 18), but only half survived the neonatal period (Table 1, row 19). EPCR^-/-^ embryos and placentae were normal in size and gross appearance at 18.5 dpc (Figure 3 D, H, I & J) but histological examination of placentae revealed thrombi in the junctional zone (Figure 3 S & T). Thus, reducing thrombin generation by inactivating FVIII was also effective in preventing early death of EPCR-deficient embryos, but insufficient in preventing late placental thrombosis and high neonatal lethality.

Nonetheless, FVIII inactivation significantly improved survival of EPCR-deficient embryos (Table 1, compare rows 1 & 18, P<0.0006, χ^2^ test of independence) and pups (Table 1, compare rows 2 & 19, P<0.037, χ^2^ test of independence). The results were comparable to Par4 inactivation which also significantly improved survival of EPCR-deficient pups (Table 1, compare rows 2 & 15, P<0.007, χ^2^ test of independence), though about half continued to die as neonates. Thus, inhibiting maternal platelet activation or reducing thrombin generation each provided significant, but partial protection, from neonatal death.

### Platelets cause placental failure independent of thrombin generation

Par4 activated platelets enhance thrombin generation. Given the similar protective effect of FVIII and Par4 inhibition, we asked whether protection from early placental failure seen in the absence of platelet receptors Par4, Par3 or αIIb might reflect their reduced ability to participate in thrombin generation. To address this question, we performed thrombin generation assays with platelet rich plasma (PRP) from Par4^-/-^, Par3^-/-^ or αIIb^-/-^ mice using recombinant TF as a trigger (Figure 4). As expected, Par4^-/-^ PRP showed reduced thrombin peak height (45.5 ± 20 nmol Par4^-/-^ vs 194.6 ± 29.2 nmol WT controls; P << 0.0001) and reduced total thrombin generation (1554.3 ± 543.2 nmol Par4^-/-^ vs 2746.9 ± 594.2 nmol WT controls; P << 0.0001), compared to wild type control. We found that Par3^-/-^ PRP also has markedly reduced thrombin peak height (53.2 ± 20.5 nmol; P << 0.0001) and reduced total thrombin generation (1300.6 ± 370.0 nmol; P << 0.0001) compared to wild type control. In contrast, the absence of αIIb did not reduce TF-induced thrombin generation by PRP *in vitro*. Rather, thrombin peak height and total thrombin generation were increased in αIIb^-/-^ PRP compared to wild type controls (peak height 289.5 ± 42.6 nmol αIIb^-/-^; P << 0.0001 and total thrombin generation 3304.9 ± 412.5 nmol αIIb^-/-^; P = 0.0003). Thus, the benefit observed with inactivation of platelet Par4 or Par3 is unlikely to be due to reduced platelet participation in thrombin generation, but rather involves other pathogenic events downstream of platelet activation.

**Figure 4:**
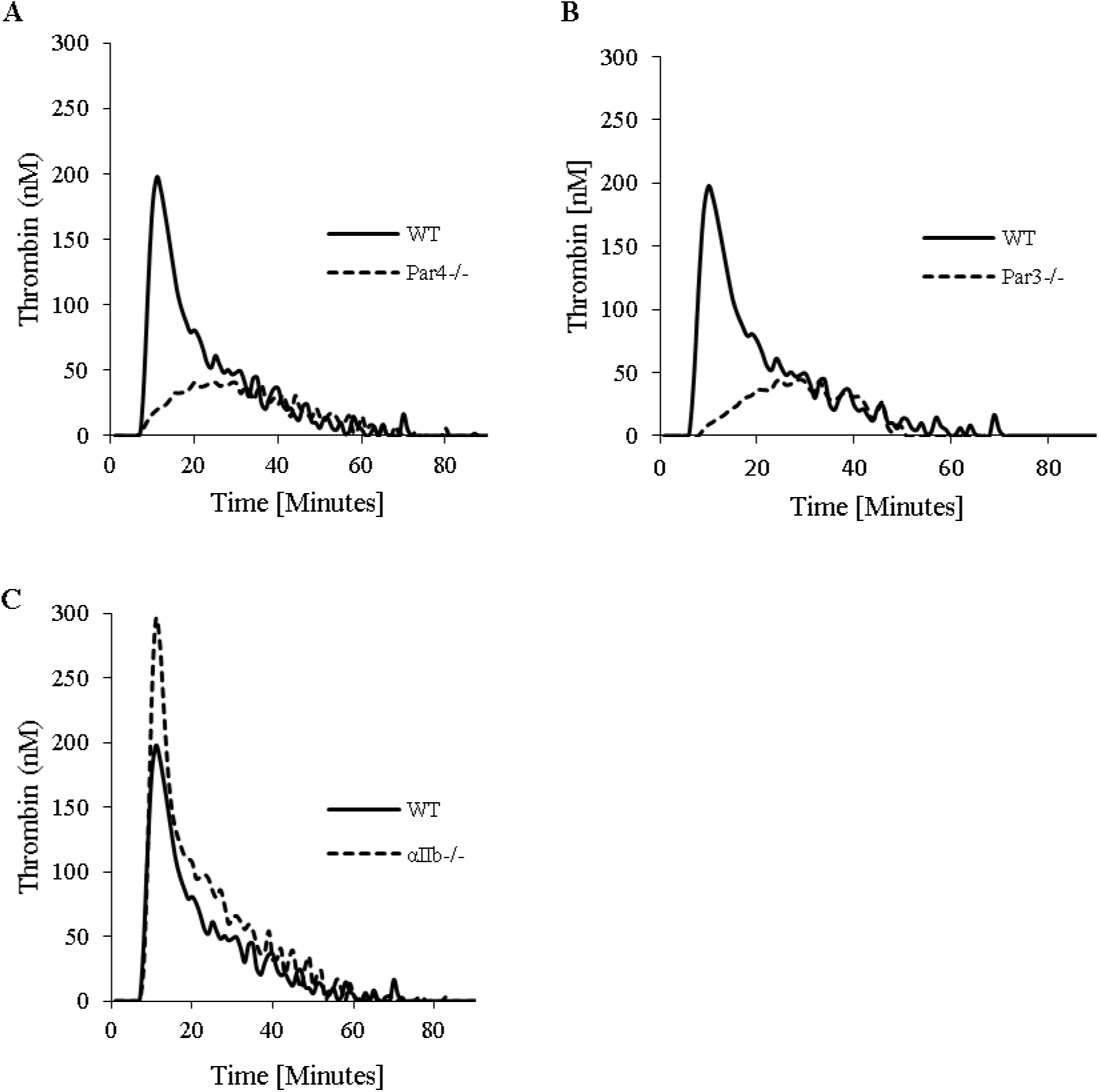
Thrombin generation triggered with rTF was significantly reduced in Par4^-/-^ (A) and Par3^-/-^ (**B**) but was enhanced in αIIb^-/-^ PRP (**C**) PRP. The assays were performed as described in the methods section.

### Maternal EPCR deficiency causes severe pregnancy complications

Adult EPCR-deficient mice generated by maintaining placental expression (EPCR^δ/δ^) exhibit plasma markers of thrombophilia without overt thrombosis ^16^. They can serve as a useful model to investigate whether maternal thrombophilia predisposes pregnancies to placenta-mediated complications. In the course of our studies, we discovered severe pregnancy complications in pregnancies of FVIII^-/-^ EPCR^-/-^ females sired by wild type males. Two of the four pregnancies that we analyzed exhibited mid-gestational vaginal bleeding. One female died towards term gestation (19.5 dpc) and three were moribund. Surgical evaluation revealed retroplacental bleeding, blanched organs and undelivered live pups. It was unclear if this was a phenomenon of concomitant EPCR and FVIII deficiencies. To address this further, we evaluated pregnancy outcome of EPCR-deficient females that were generated by maintaining EPCR expression in the placenta (EPCR^δ/δ^). We analyzed 18 pregnancies of 12 dams sired by wild type males. Of these, 8 pregnancies (44%) resulted in maternal death or moribund mother requiring euthanasia. One maternal death occurred on 13.5 dpc and two on 15.5 dpc. In the remaining 5 cases the mother was moribund past the expected delivery date; 4 of these were associated with dead or live pups that had not been delivered. In addition, 2/18 pregnancies resulted in stillborn pups, 1/18 in pregnancy loss, and 8/18 showed mid-gestational vaginal bleeding. Surgical evaluation of pregnant females at mid-gestation revealed retroplacental bleeding. Surgeries at term revealed more extensive retroplacental hemorrhaging paired with blanched appearance of maternal organs. Upon histological evaluation, the placentae appeared normally developed but blood clots were observed in the decidua basalis, both at mid and late gestation (Figure 5 A-D, I, J). In contrast, the placental layers (giant trophoblast cells, junctional zone and labyrinth) were comparable to wild type controls and remarkably devoid of thrombotic pathology (Figure 5 E-H, K, L). Thus, genetic inactivation of EPCR in female mice results in decidual thrombosis, retroplacental hemorrhaging and vaginal bleeding at mid-gestation, complications of parturition and frequent maternal and/or fetal death. Overall, these findings are consistent with placental abruption and consumptive coagulopathy, suggesting that maternal death may be caused by hemorrhagic shock. Immunohistochemical staining of placentae revealed infiltration of neutrophils, release of myeloperoxidase and positive staining for citrullinated histone 3, indicative of neutrophil extracellular trap formation (Figure 5 M-R). We evaluated pregnancy outcome of EPCR-deficient mice with a concomitant deficiency of Par4. In striking contrast, five pregnancies of Par4^-/-^ EPCR^-/-^ dams sired by wild type males resulted in uneventful pregnancies with normal outcomes. These data demonstrate an essential role of maternal EPCR in the maintenance of pregnancy. Inactivation of Par4, but not clotting factor VIII, prevented maternal death and restored normal pregnancy outcomes.

**Figure 5:**
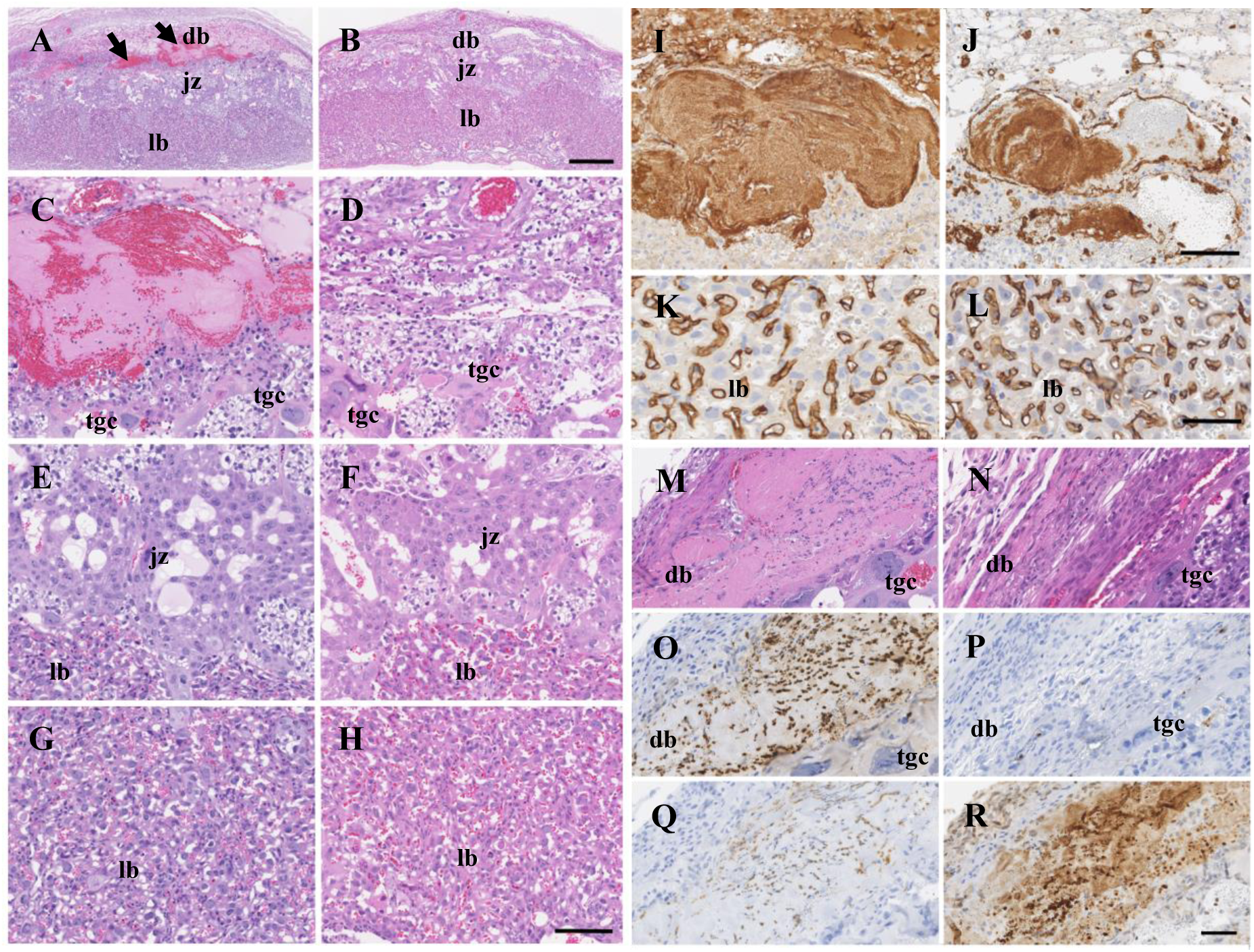
EPCR-deficient (EPCR^δ/δ^) dams exhibit frequent thrombosis in the decidua basalis but not in the placenta. Hematoxylin-eosin stained histological sections of 15.5 dpc placentae harvested from wild type intercrosses (**B**) and from pregnancies of EPCR^δ/δ^ dams sired by wild type males (**A**) are shown. Arrows in A point to regions of decidual thrombosis. Magnified views of decidua basalis (**C, D**), junctional zones (**E, F**) and labyrinth regions (**G, H**) of placentae in A and B are shown. Decidual thrombosis was confirmed by immunostaining with fibrin(ogen) (**I**) and CD31 (**J**) antibodies. CD31 immunostaining confirmed normal elaboration of placental labyrinth (**K**) comparable to placenta from control wild type pregnancies (**L**). Decidual thrombosis was also readily seen in hematoxylin-eosin stained histological sections of 12.5 dpc placentae from pregnancies of EPCR^δ/δ^ dams (**M**) but not in gestational age-matched wild type pregnancies (**N**). Strong immunostaining was observed with myeloperoxidase antibodies (**O**; myeloperoxidase stained placenta from control pregnancy shown in **P**), neutrophil marker Ly6G (**Q**) and antibodies against citrullinated histone 3 (**R**). db: decidua basalis; jz: junctional zone; lb: labyrinth; tgc: trophoblast giant cells. Scale bars=500µm (A and B) or 100 µm C-H, I-J), 50μm (K-R).

## DISCUSSION

In this work, we used genetic strategies to first examine whether thrombin generation and maternal platelet activation play key roles in placental failure due to fetal thrombophilia. We demonstrate that thrombin-mediated activation of maternal platelets causes the early placental failure and mid-gestational death of EPCR-deficient murine embryos. Inactivation of FVIII, maternal Par4, Par3 or αIIb are each equally effective in allowing placental development and survival of EPCR-deficient embryos. Once past mid-gestation, lack of extraembryonic EPCR results in thrombosis specific to maternal venous sinuses in the junctional zone of the placenta. These are continuous with decidual and uterine veins and are analogous to extravascular trophoblast lined venous channels in the basal plate of human placenta. Our study identifies these venous channels as potential sites of thrombotic pathology in human pregnancies. It was recently shown that human extravillous trophoblasts not only invade uterine arteries, but also line decidual and uterine veins in early pregnancy ^33,34^, extending potential sites of pathology to these regions. Inhibition of thrombin-mediated platelet activation or reduction of thrombin generation alone were each insufficient to fully overcome late gestation placental thrombosis. Despite these interventions, placental EPCR deficiency resulted in high neonatal lethality not observed in EPCR-deficient mice with intact expression in the placenta. Our observations indicate gestation age-specific differences in fetal thrombophilia-associated placental pathology and its potential response to treatment. They lend support to the notion that placental pathology is an important contributor to neonatal death ^35,36^. Notably, both anti-thrombotic approaches improved embryonic and neonatal survival compared to no intervention.

Our study has uncovered a critical requirement of maternal EPCR in the maintenance of pregnancy. We demonstrate that decidual thrombosis, retroplacental hemorrhaging, vaginal bleeding, and consumptive coagulopathy are components of pathogenic mechanisms resulting in adverse fetal outcomes and pregnancy-induced death of EPCR-deficient dams. These findings directly implicate coagulation abnormalities in placental abruption in human pregnancy. Importantly, we demonstrate that inactivation of Par4 overcomes these severe pregnancy complications and prevents maternal death of EPCR-deficient mice. In contrast, reduced thrombin generation achieved by inactivation of VIII is ineffective in this scenario. Our findings lend credence to the notion that thrombotic processes mediate adverse pregnancy outcomes associated with maternal or fetal thrombophilia and draw attention to the potential for therapeutic inhibition of thrombin-mediated platelet activation in a subset of pregnancy complications. Conversely, our results indicate that polymorphisms associated with increased Par4 reactivity ^37,38^ should be evaluated for a potential association with gestational complications, particularly in populations of African ancestral descent where both conditions are more prevalent. The new murine model of pregnancy-induced maternal death provides an opportunity to elucidate the mechanism of maternal thrombophilia-associated pregnancy complications and evaluate therapeutic potential of anti-Par4 and other anti-platelet agents in prevention of adverse pregnancy outcomes.

## Acknowledgments

The authors thank Michelle Bordas and Katelyn Storage for help with maintaining the murine colony and genotyping, Gwendolyn Werra for immunostaining of murine placenta with platelet antibodies, Kirkwood Pritchard for MPO and Cit H3 antibodies, the CRI Histology and Imaging cores for expert services, and Andrew Webb, Valerie Salato and Ann Diamond for administrative support.

## Author contributions

M.M.C., Q.Y., A.S., D.T.M., M.Z. and K.L.C. performed experiments. J.A.J. examined and verified pathology observations. M.M.C. and R.S. analyzed data and prepared figures and tables. R.S. formulated the study, designed methodology, provided supervision and wrote the manuscript. M.M.C and J.A.J. critically reviewed the manuscript.

## SUPPLEMENTARY TEXT

### Note 1

Aspirin inhibits Cox-1 and Cox-2 enzymes, and consequently, the conversion of arachidonic acid into thromboxane A2, a platelet agonist. Pregnant dams from EPCR^+/-^ intercrosses were fed 40 or 80mg/kg/day of Aspirin daily by oral gavage, starting at 5.5 days post coitum (dpc). Aspirin treatment of pregnant dams effectively eliminated arachidonic acid-induced platelet aggregation (Figure S1 L & M) and resulted in two severely growth restricted EPCR-deficient embryos out of 28 live embryos analyzed at 11.5/12.5 dpc (12 were aborted). The placentae of these embryos showed thickened maternal decidua and giant cells, but no labyrinth was elaborated (Figure S1 A-E). Thus, aspirin treatment prolonged survival of a small fraction of EPCR-deficient embryos but did not restore normal placental development.

### Note 2

We examined whether depletion of platelets allows development of EPCR^-/-^ placentae past 10.5 dpc. Plugged females from EPCR^+/-^ intercrosses were injected at 7.5 dpc with platelet depleting antibodies directed against GP1b. Treatment results in transient thrombocytopenia in the pregnant dam that lasts for 4 days after treatment (Figure S1 J & K). Analysis of two pregnancies at 11.5 dpc showed 2/16 embryos to be EPCR^-/-^. In another treated pregnancy analyzed at 15.5 dpc, 1 of 6 embryos was EPCR^-/-^. In all three pregnancies, EPCR-deficient embryos and placentae were normal in size, appearance, and histological features (Figure S1 F-I). Thus, treatment of the mother with platelet depleting antibodies restored placental development and improved survival of EPCR-deficient embryos.

### Note 3

To be able to examine if LMWH restores development of EPCR^-/-^ placentae, we repeated the previously published experiment where pregnant dams from EPCR^+/-^ intercrosses were treated with LMWH ^16^. To achieve continuous anticoagulation, we implanted pregnant females with subcutaneous osmotic pumps that delivered LMWH at 25μg/hour from 5.5 dpc till the time that pregnancies were analyzed. Treatment resulted in 3 EPCR^-/-^ out of 19 live embryos analyzed at 11.5 dpc. All three embryos were smaller, and one was severely growth restricted. All three placentae showed marked reduction in placental labyrinth formation (Figure S2). Treatment resulted in significant anticoagulation measured as 0.3 to 0.5 IU of FXa inhibitory activity per mL of maternal plasma. Thus, normal placental development could not be restored even with continuous anticoagulation with LMWH.

## MATERIALS AND METHODS

### Mice

Par4 (MMRRC stock number: 15231), Par3 (MMRRC stock number: 15232), αIIb, FVIII, EPCR loxP and Meox2Cre (Jackson laboratory, stock number 003755) mice have been previously described ^30, 39-42, 16, 43^. All mouse strains were maintained on C57BL/6 genetic background. Double mutants were generated by breeding and identified by polymerase chain reaction-based genotyping on tissue obtained by tail or ear biopsy. Where noted, the PCR primers recommended by the Mutant Mouse Regional Resource Center (MMRRC) or Jackson laboratory were used. For other strains, PCR primers described in the original referenced publications were used. All animal experiments were conducted following standards and procedures approved by the Animal Care and Use Committee of the Medical College of Wisconsin.

### Analysis of pregnancies

Paired mice were checked every morning for presence of vaginal plug as evidence of coitum. On the day of the plug, 12 Noon was counted as 0.5 dpc. Uteri were surgically removed and dissected in phosphate buffered saline. Embryos and placentae were examined and photographed with a Nikon SMZ1000 Zoom stereomicroscope (Nikon, Melville, NY) equipped with a high resolution, 5 megapixel digital camera and NIS Elements F4.3 imaging software. Live embryos were identified by the presence of heartbeats, breathing or limb movements. Embryos and placentae were observed for any phenotypic abnormalities. DNA was prepared from the yolk sacs or tail biopsies of embryo. Presence or absence of genes of interest were determined by polymerase chain reaction using gene specific primers. Embryos found at advanced stages of resorption were not genotyped. The placentae were marked with ink to identify the center of the disc in histological sections. To estimate relative sizes, the cross-sectional area of placental discs and lengthwise cross-sectional area of the embryos was measured with image J (version 1.52a; National Institutes of Health, Bethesda, MD) from images of whole mounts as shown in Figure 3 A-D.

### Histology and Immunohistochemistry

For most experiments, tissues were fixed in zinc formalin for 72 hours and transferred to 70% alcohol for future processing. Fixed tissues were later embedded in paraffin, sectioned and stained with hematoxylin-eosin using standard protocols. For immunostaining, antigen retrieval was performed in BOND epitope retrieval solution 1 (pH 6.0) or solution 2 (pH 9.0) (Leica Biosystems, Buffalo Grove, IL). EPCR immunostaining was done with anti-EPCR primary antibodies at a dilution of 1:200 (Cat # AF2749, R&D systems, Minneapolis, MN) and biotinylated rabbit anti-goat secondary antibodies at a dilution of 1:500 (Cat # BA-5000, Vector Labs, Burlingame, CA). Cytokeratin, CD31 and fibrin(ogen) immunostaining was done with 1:500 dilution of anti-cytokeratin (Cat # Z0622, Dako, Santa Clara, CA) or 1:100 dilution of anti-CD31 (Cat # AF3628, R&D systems, Minneapolis, MN) or 1:2000 anti-fibrinogen (Cat # A0080, Dako, Santa Clara, CA) primary antibodies and 1:500 dilution of biotinylated donkey anti-rabbit secondary antibodies (Cat # 711-066-152, Jackson ImmunoResearch laboratories, West Grove, PA). Neutrophils were identified with anti-Ly6G primary at 1:75 (Cat # 14-5931-85, eBioscience, San Diego, CA) and biotinylated goat anti-rat IgG secondary antibodies at a dilution of 1:300 (Cat # BA-9401, Vector labs, Burlingame, CA). In all of the above immunostaining protocols, streptavidin/HRP was used at a dilution of 1:300 for 15 minutes (Cat # P039701-2, Dako, Santa Clara, CA) and DAB+ kit (Cat # K346811-2, Dako, Santa Clara, CA) was used for detection. Immunostaining for myeloperoxidase and citrullinated histone 3 was done with anti-MPO or anti-CitH3 primary antibodies (Cat # ab208670, Abcam, Cambridge, MA; Cat # ab5103, Abcam, Cambridge, MA) at 1:500 dilution each and developed with the BOND polymer refine detection kit (Cat # DS9800, Leica Biosystems, Buffalo Grove, IL). Stained histological sections were photographed and viewed using a Nanozoomer HT 2.0 slide scanner and NDP view-imaging software (Hamamatsu, Japan). For some experiments, placentae were embedded and flash frozen in OCT, fixed in acetone after sectioning, and immunostained with rabbit anti-Cow Cytokeratin (Cat# Z0622, Dako, Santa Clara, CA) and rat anti-mouse GPIbβ (Cat # M050-0, Emfret Analytics, Germany) primary antibodies at 1:100 dilution each. Primary cytokeratin antibodies were visualized with goat anti-rabbit IgG Alexa Fluor 633 (Cat # A-21071, Invitrogen, Carlsbad, CA). Primary GPIbβ antibodies were visualized with 4plus Biotinylated Goat Anti-Rat IgG (Cat # GR607, Biocare Medical) and Streptavidin 488 (Cat # S11223, Invitrogen, Carlsbad, CA). Images were taken on a Zeiss LSM510 confocal system (Zeiss International, Germany).

### Drug and antibody treatments

To achieve continuous anticoagulation with LMWH, pregnant dams were implanted with subcutaneous osmotic pumps (Model 1002, Durect Corp, Cupertino, CA) filled with 100μg/μL LMWH (enoxaparin, brand name Lovenox; Aventis Pharmaceuticals Inc, Bridgewater, NJ) with release rates of 0.25 μL/hour. Treatment was initiated at 5.5 dpc and continued until the time of analysis at 11.5 dpc. Anticoagulation was measured in terms of Xa inhibitory activity with Coatest heparin (Chromogenix, Lexington, MA) in plasma. Plasma was prepared from citrated whole blood collected from the inferior vena cava at the time of analysis of pregnancies. To deplete platelets, pregnant dams were intravenously injected with anti-GPIbα antibodies (R300, Emfret analytics GmbH & Co.KG, Wurzburg, Germany) at 7.5 dpc at a dose of 4 µg/g body weight. Platelet depletion was confirmed one hour after treatment by flow cytometry on 5μL whole blood collected from the tail vein, diluted in citrated Tyrode’s buffer and stained with anti-CD41 eFlour 450-labelled antibodies (Cat # 48-0411-80, eBioscience, San Diego, CA) in the dark for 15 minutes. In a separate set of experiments, aspirin (acetylsalicylic acid, Cat # A5376, Sigma Aldrich, St. Louis, MO) was administered daily at 40 or 80 mg/kg via oral gavage from 5.5 dpc to the day of analysis of pregnancies. At the time of analysis, citrated whole blood collected from the inferior vena cava was used to prepare washed platelets. Whole blood was centrifuged at 150 × g for 5 minutes and the supernatant was collected and centrifuged again at 800 × g for 8 minutes to sediment platelets. Platelets were washed with Tyrode’s buffer containing 2.5 mg/mL BSA and 1mg/mL glucose, and centrifuged at 800 × g for 8 minutes and finally re-suspended in the same buffer. All centrifugation steps were done at room temperature and without brakes. To inhibit platelet activation, Prostaglandin E1 (PGE1) (Cat # P5515, Sigma Aldrich, St. Louis, MO) was added prior to each centrifugation step at a final concentration of 50ng/mL. To inactivate any residual PGE1, platelet suspension was rested for 30 minutes at room temperature prior to the aggregation experiment. Prior to measuring aggregation response, calcium was added to the platelet suspension at a final concentration of 1mM. The aggregation response of platelets was measured following stimulation with arachidonic acid (Cat. No.: P/N 390, Chrono-log, Havertown, PA) or U46619 (Enzo Life Sciences, Farmingdale, NY) using the Chrono-log 700 instrument (Chrono-log, Havertown, PA).

### Thrombin generation assay

To prepare Platelet Rich Plasma (PRP), citrated whole blood collected from the inferior vena cava was centrifuged at 150 × g for 5 minutes without brakes. The supernatant was collected, and platelet counts were measured on a Scil Vet ABC Plus Hematology Analyzer (Scil animal care company, Ontario, Canada). Platelet counts were adjusted to 600×10^3^ platelets/µL using Platelet Poor Plasma (PPP) prepared from the same whole blood sample using a second spin at 800 × g for 8 minutes. *In vitro* thrombin generation assays were conducted using a modified Technothrombin® Thrombin Generation Assay (Cat # 5006010, DiaPharma Group, West Chester Township, OH). Thrombin generation was initiated by the addition of rTF [1.0pM] (RecombiPlasTin 2G, Instrumentation Laboratory, Bedford, MA). Fluorescence was immediately measured with a SpectraMax Gemini EM fluorimeter and SoftMax pro software (Molecular Devices, San Jose, CA) at 355nm/460 nm (excitation/emission) for two hours at 37°C with measurements collected at one-minute intervals. Raw fluorescence data was analyzed using Technothrombin TGA evaluation software.

### Statistical analysis

The χ2 goodness-of-fit (GOF) test was used to determine deviation from expected Mendelian proportions. The χ2 test of independence was used for comparing two separate treatment groups. Exact binomial 95% confidence intervals (CI) were computed where appropriate. The Student’s t test (2 tailed with unequal variance) was used for comparing embryo and placental sizes between groups and for comparing fluorescence values in thrombin generation assays. For all experiments, P < 0.05 was considered significant.

**Figure S1:**
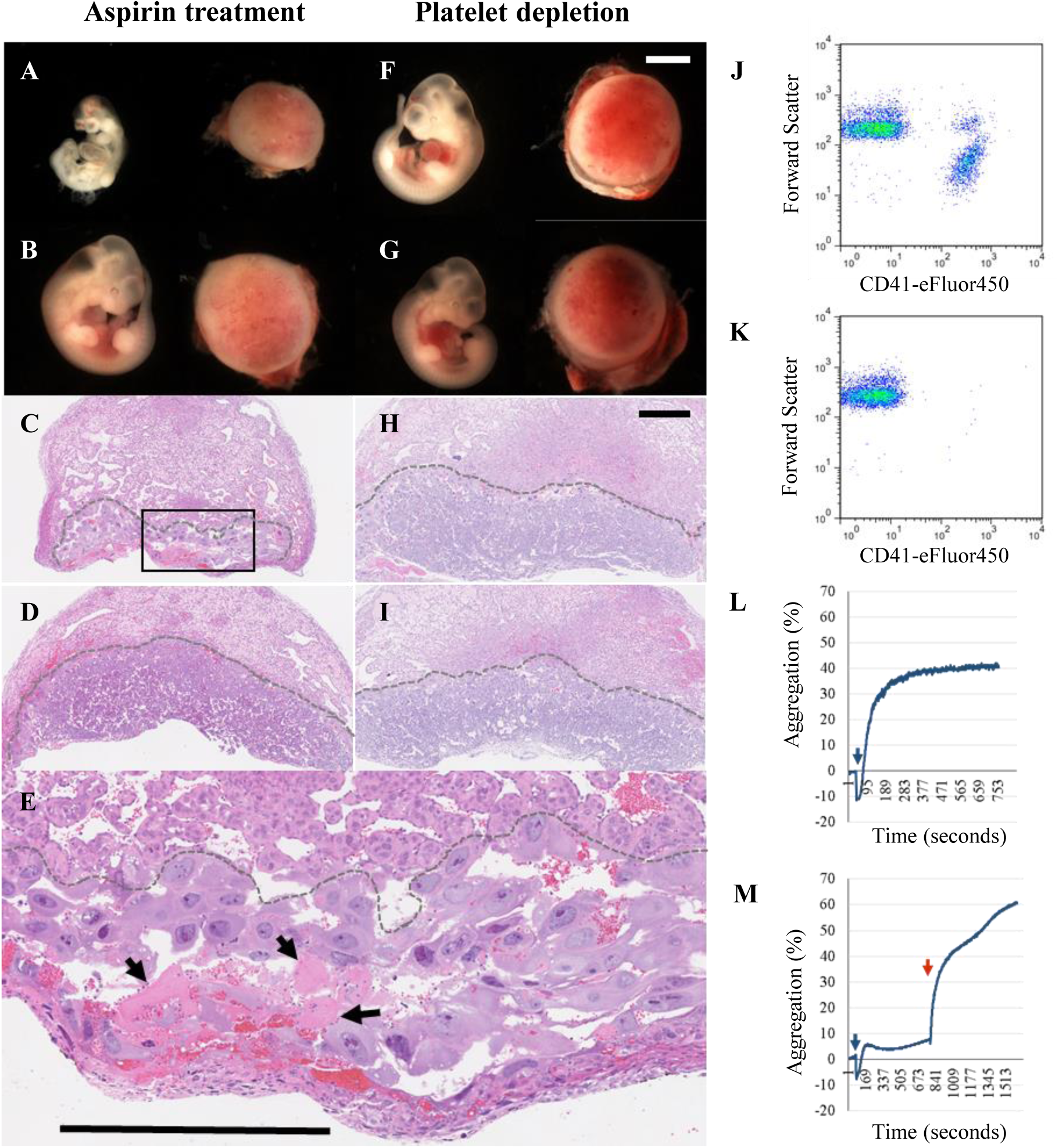
Placental development of EPCR-deficient embryos is restored in the absence of platelets but not by treatment of the mother with aspirin. Treatment with aspirin prolonged survival of EPCR^-/-^ embryos, but they were severely growth restricted (**A**) compared to littermate EPCR^+/+^ controls (**B**). EPCR^-/-^ placentae (**C**, boxed area enlarged in **E**) did not form a labyrinth and continued to exhibit blood clots (marked by arrows in E). Labyrinth was elaborated in placentae of littermate EPCR^+/+^ controls (**D**). Antibody-mediated depletion of maternal platelets restored development of EPCR^-/-^ embryos and placentae (**F**) comparable to littermate EPCR^+/+^ controls (**G**). EPCR^-/-^ placentae (**H**) showed labyrinth formation comparable to EPCR^+/+^ controls (**I**). Scale bars represent 2mm (whole mounts) or 500μm (placental sections). J and K are scatter plots of flow cytometry data points. Platelets identified by CD41 staining in whole blood from untreated mice (**J**) are absent in whole blood of pregnant females treated with platelet depleting antibodies (**K**). L and M are aggregometer tracings showing the effect of aspirin treatment on maternal platelet aggregation. Addition of arachidonic acid (blue arrows) did not stimulate aggregation of platelets obtained from aspirin treated pregnancies (**M**) but was effective in control samples from untreated mice (**L**). U46619 (thromboxane A2 receptor agonist) was added (red arrow) as a positive control to verify that the platelets aggregated in response to other agonists.

**Figure S2:**
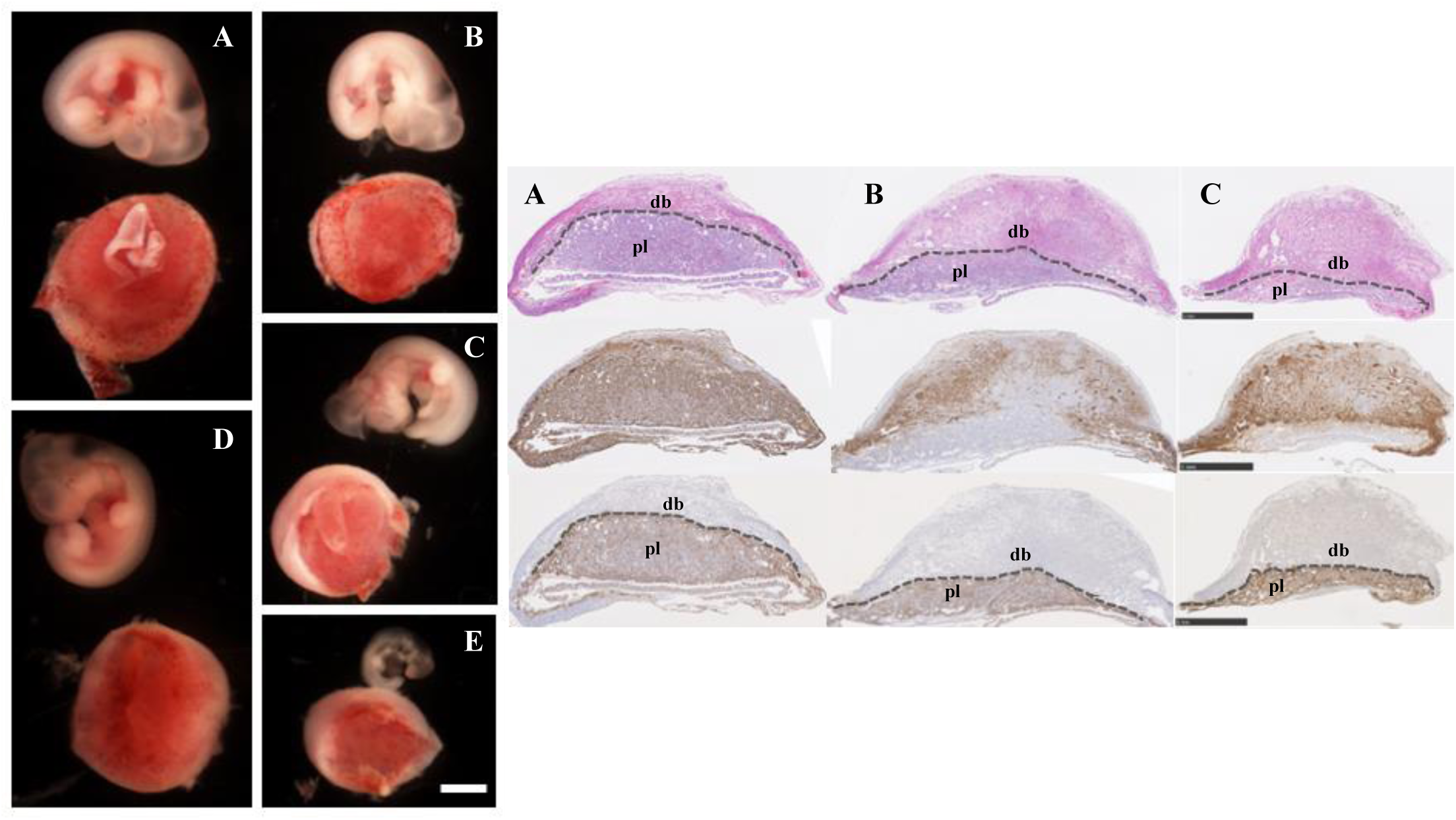
Continuous anticoagulation of the mother with LMWH did not restore normal placental development of EPCR-deficient embryos. Whole mount of EPCR^-/-^ (**B, C, E**) and littermate EPCR^+/+^ (**A, D**) embryos and placentae from LMWH treated pregnancies are shown in the left panel and placental sections corresponding to A, B and C stained with hematoxylin-eosin (top row) or immunostained with EPCR (middle row) or cytokeratin antibodies (bottom row) are shown in the right panel. Placental regions identified with hematoxylin-eosin staining or cytokeratin expression of trophoblast cells are delineated with dashed lines. EPCR is not expressed in placentae corresponding to EPCR^-/-^ embryos but continues to be expressed in maternal decidua. db: decidua basalis, pl: placenta. Scale bars represent 2mm (whole mounts) or 1mm (placental sections).

